# What the hippocampus tells the HPA axis: Hippocampal output attenuates acute stress responses via disynaptic inhibition of CRF+ PVN neurons

**DOI:** 10.1101/2022.04.14.488387

**Authors:** Anthony B. Cole, Kristen Montgomery, Tracy L. Bale, Scott M. Thompson

## Abstract

The hippocampus exerts inhibitory feedback on the release of glucocorticoids. Because the major hippocampal efferent projections are excitatory, it has been hypothesized that this feedback inhibition is mediated by populations of inhibitory neurons in the hypothalamus or elsewhere. These regions would be excited by hippocampal efferents and project to corticotropin-releasing factor (CRF) cells in the paraventricular nucleus of the hypothalamus (PVN). A direct demonstration of the synaptic responses elicited by hippocampal outputs in PVN cells or upstream GABAergic interneurons has not been provided previously. Here, we used viral vectors to express channelrhodopsin (ChR) and enhanced yellow fluorescent protein (EYFP) in pyramidal cells in the ventral hippocampus (vHip) in mice expressing tdTomato in GABA-or CRF-expressing neurons. We observed dense innervation of the bed nucleus of the stria terminalis (BNST) by labelled vHip axons and sparse labeling within the PVN. Using whole-cell voltage-clamp recording in parasagittal brain slices containing the BNST and PVN, photostimulation of vHip terminals elicited monosynaptic excitatory postsynaptic currents (EPSCs) and disynaptic inhibitory postsynaptic potentials (IPSCs) in both CRF+ and GAD+ cells. The balance between synaptic excitation and inhibition were maintained in CRF+ cells during 20 Hz stimulus trains. Photostimulation of hippocampal afferents to the BNST and PVN *in vivo* inhibited the rise in blood glucocorticoid levels produced by acute restraint stress. We thus provide functional evidence that hippocampal output to the BNST results in a net inhibition of the hypothalamic-pituitary axis, gaining further mechanistic insights into this process using methods with enhanced spatial and temporal resolution.

## 1. Introduction

The corticosteroid neuroendocrine response is controlled by the hypothalamic-pituitary-adrenal (HPA) axis. (for review ^1–4^) Corticotropin-releasing factor (CRF)-releasing neurons in the paraventricular nucleus of the hypothalamus (PVN) are the final common site upon which central projections from stress-sensitive brain regions converge to modulate HPA axis activity. CRF reaches corticotropes in the pituitary gland through the hypophyseal-portal circulation and stimulates the release of adrenocorticotropic hormone (ACTH), which then enters systemic circulation. Cells within the adrenal cortex are stimulated by ACTH binding to the Melanocortin receptor type 2 to synthesize and release glucocorticoids into circulation. The end product of HPA activation is the rapid production of glucocorticoids that have powerful and systemic effects on numerous systems throughout the body. The production of glucocorticoids therefore also requires tight regulation, typically thought to occur through negative feedback via glucocorticoid receptor (GR) and mineralocorticoid receptor (MR)-expressing cells. The canonical actions of these intracellular receptors act via relatively slow genomic mechanisms. During an acute stress event in a healthy functioning HPA axis, peak glucocorticoid levels rise rapidly, peaking within 30 minutes of stress onset, and return back to baseline within 120 minutes via negative feedback ^1^. Numerous clinical manifestations can develop when glucocorticoids are chronically produced or inadequately controlled, including central fat deposits, hair loss, loss of bone density, and psychiatric manifestations ^5,6^. Both hypo-and hyper-activation of the HPA axis is also a consistent finding in PTSD and major depressive disorder, and has been suggested to be an etiological factor in their genesis ^7–11^.

GR and MRs are differentially expressed throughout the brain but are concentrated in regions thought to be responsible for mediating responses to stressful conditions. The ventral hippocampus (vHip) carries affective information, displays high levels of MR/GR, and has been implicated in HPA regulation ^12,13^. In support of this hypothesis, lesioning the vHip or major hippocampal efferent tracts leads to alterations in the responses to acute stressors ^14,15^ and an inability of exogenous corticosteroids to suppress HPA activation ^16^. Genetic deletion of GRs in cortical and hippocampal pyramidal cells also results in an elevation of circulating glucocorticoids ^14^.

The output of hippocampal pyramidal cells is glutamatergic, and should excite CRF cells in the PVN if they are synaptically connected. It is suggested that an interposed population of GABAergic cells reverses this excitatory drive to inhibit the HPA axis. Electrophysiological studies have identified several candidate brain regions with a high density of inhibitory neurons that project to the PVN ^17^, but to our knowledge there is no physiological evidence demonstrating that these cells are both excited by hippocampal output and inhibit PVN cells. The bed nucleus of the stria terminalis (BNST) is one such candidate region ^18–22^. It has dense expression of GAD+ and CRF+ neurons and is positioned to integrate many stress-relevant signals and relay them to the PVN ^19,20,23,24^. Lesions and optogenetic manipulations within different subnuclei or subregions of the BNST have varying effects on HPA response ^25–28^. Our study uses this groundwork to guide our investigation into how exactly the hippocampus regulates the HPA axis and which intermediate brain regions may play a role.

## 2. Methods

All procedures were approved by the University of Maryland School of Medicine Institutional Animal Use and Care Committee and performed in accordance with the National Institutes of Health Guide for the Care and Use of Laboratory Animals.

### 2.1. Animals and housing

Male and female mice, 5 - 20 weeks of age, were group-housed with *ad libitum* access to water and standard rodent chow. Reporter mice were used for identification of the cell populations of interest by crossing Ai14 mice (B6;129S6-*Gt(ROSA)26Sor*^*tm14(CAG-tdTomato)Hze*^/J; Jackson Laboratory), for cre-dependent expression of tdTomato, with mice expressing cre recombinase under the control of the CRF (B6(Cg)-*Crh*^*tm1(cre)Zjh*^/J; Jackson Laboratory) or glutamic acid decarboxylase (*Gad2*^*tm2(cre)Zjh*^/J; Jackson Laboratory) promoters ^29–32^.

### 2.2. Viral injection

Stereotaxic surgery (Kopf stereotaxic apparatus) was performed on male and female mice at 5-6 weeks of age. AAV2 viral vectors were used to express channelrhodopsin and enhanced yellow fluorescent protein (EYFP)(AAV2-CAMKIIa-hChR2(H134R)-EYFP, UNC Viral Core) in Ca^2+^/calmodulin-dependent kinase expressing neurons, likely pyramidal cells ^33,34^. Viral solutions (0.5 μl, titer 1-8×10^12vg/ml) were injected bilaterally into ventral CA1 of the hippocampus (mm to Bregma: 3.3 AP, ±2.8 Lat, -3.8 D/V) at a rate of 0.1μl/minute. The syringe (Hamilton Neuros syringe, 33 gauge) was left in place for ten minutes following final injection to allow for dispersal before withdrawing the syringe.

### 2.3 Imaging

Mice were sacrificed and brain tissue was collected five weeks following injection, allowing for expression and transport of ChR to distal axon terminals. Animals were transcardially perfused using 30ml of phosphate-buffered saline (PBS, containing in mM: 137 NaCl, 2.7 KCl, 10 Na_2_HPO_4_, 1.8 KH_2_PO_4_), followed by 4% paraformaldehyde (PFA) in PBS. The brain was then extracted and mounted on to an agar base for vibratome sectioning. Brain slices for imaging were sectioned in either the coronal or parasagittal plane at a thickness of 75 μm on a Leica VT 1200S vibratome. Slices were washed in deionized water prior to slide mounting using Vectashield+DAPI. (Vector Laboratories, Burlington, CA) Images were taken on an Nikon Eclipse E600FN upright microscope (Fig S1), Nikon W1 spinning disk confocal microscope fitted with Hamamatsu sCMOS camera (Fig 1c,d,e), or Nikon Ti2-E inverted epifluorescence microscope fitted with a Spectra-X

**Figure 1.**
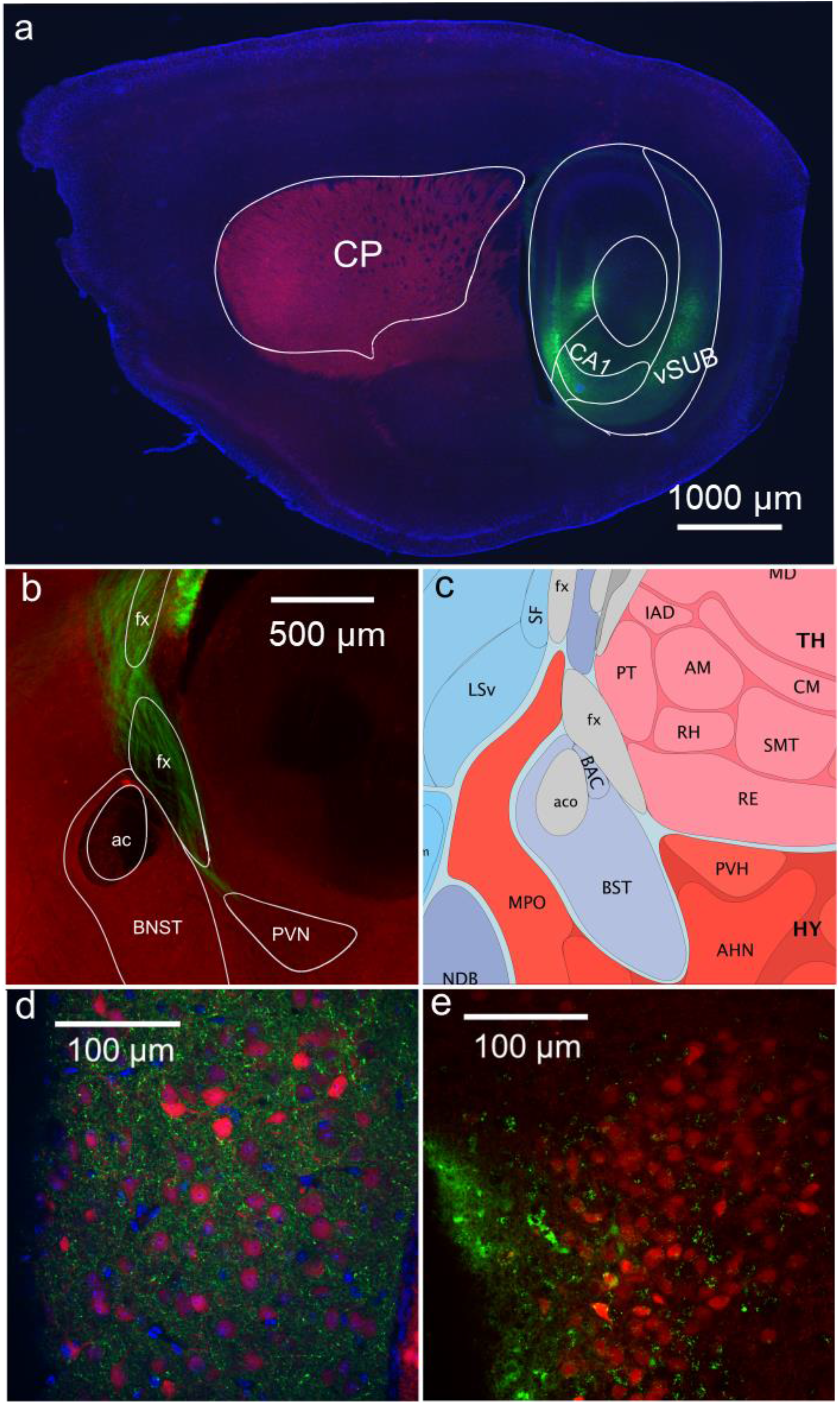
Anatomy of ventral hippocampal terminals in GAD and CRF reporter mice. **a)** EYFP (green) in glutamatergic pyramidal cells in the ventral hippocampus and subiculum using an AAV2 virus with CAMKII-promoter driven expression in a GAD-tdTomato (red) mouse brain. Section sliced in the parasagittal plane and annotated with outlines from Allen Brain Atlas (sagittal view 2/21). **b)** Strong expression of EYFP (green) was seen in axons coursing through the fornix (fx), both anterior and posterior of the anterior commissure (ac), in a GAD-tdTomato (red) mouse. **c)** Diagram of nuclei in parasagittal sections of this region (Allen Mouse Brain Atlas and Allen Reference Atlas - Mouse Brain, sagittal view 19/21 image credit: Allen Institute for Brain Science https://atlas.brain-map.org/atlas?atlas=2#atlas=2&plate=100883858). d) Numerous hippocampal terminals (green) in the medial regions of the BNST in close proximity to GAD+ neurons (red). **e)** Sparse hippocampal terminals (green) near CRF+ neurons in the PVN in a coronal sections from a CRF-tdTomato (red) mouse. DAPI counterstain (blue) in all sections.

7 channel LED light engine (Lumencor, Beaverton, OR) (Fig1a,b), as noted. Brain regions were referenced from the Mouse Brain Atlas and outlines from the atlas were used in the creation of anatomical images. (Allen Sagittal Images 19/21, Coronal Images 57/132, 59/132)^35–39^ BNST subnuclei were identified using landmarks outlined in Lebow and Chen (2016) ^24^.

### 2.4 Electrophysiology of hippocampal projections in vitro

Five weeks after viral delivery of ChR-YFP, mice were euthanized using isoflurane overdose and then perfused with critical recovery solution (CRS; in mM N-methyl D-glucamine 92, KCl 2.5, NaH_2_PO_4_ 30, HEPES 20, glucose 25, CaCl_2_ 0.5, MgCl_2_ 10) prior to decapitation. ^40^ The brain was removed and mounted in ice-cold CRS during slicing. Coronal and parasagittal sections (300 μm thickness) were obtained, rested for 12 minutes, and then transferred to HEPES-buffered holding solution (NaCl 92, KCl 2.5, NaH_2_PO_4_ 1.25, NaHCO_3_ 30, HEPES 20, glucose 25, CaCl_2_ 2, MgCl_2_ 2) for 1 hour prior to recording. Unless otherwise stated, all data shown were from slices taken in the parasagittal plane. Patch clamp recording was performed using Clampex 10.7 software, Digidata 1440 digitizer, and an Axopatch 200B amplifier (Molecular Devices, San Jose, CA). Pipettes were pulled to 4-10 mOhms using a Sutter Instruments P-87 micropipette puller and filled with a cesium methylsulphate-based pipette solution (concentrations in mM, CsCH_3_SO_4_ 135, MgCl_6_-H2O 2, HEPES 10, Mg-ATP 4, Na_2_-GTP 0.3, Na_2_-phosphocreatine 10, K_4_-BAPTA 10). Slices were recorded in whole-cell voltage-clamp configuration in artificial cerebrospinal fluid (ACSF; in mM, NaCl 124, KCl 3, NaH_2_PO_4_ 1.25, NaHCO_3_ 26, glucose 20, MgCl_2_ 1.5, CaCl_2_ 2.5) bubbled with carbogen (95% CO_2_,

5% O_2_).

Optically evoked synaptic currents were elicited with a Prismatix BlueLED light source (Southfield, MI) (460nm, 1ms pulse duration 5-10mW) delivered by an optical fiber with a 1000μm diameter core (Thor Labs, Newton, NJ). The fiber was positioned with a micromanipulator to areas of dense EYFP expression near the junction of the fornix and BNST, as visualized with epifluorescence on the recording setup. Electrically evoked potentials were driven by direct stimulation of the fornix using a concentric bipolar electrode. Square current pulses of 1 ms duration and an intensity of 0.1-10mA (World Precision Instruments Isostim A320, Sarasota, FL) were delivered either as single pulses at 0.1 Hz or as a train of pulses at 20 Hz, with 1 minute between stimulus trains. The CsMeS-based internal pipette solution blocks K channels and facilitates clamping to depolarized holding potentials. Conductances were calculated using the amplitude of the synaptic current divided by the driving force for the channels of interest (AMPAR-mediated current reversal potential = 0mV, GABA_A_R-mediated current reversal potential = -60mV).

### 2.5. Stimulation of hippocampal projections in vivo

Optical fibers were constructed using a modified protocol as described by Sparta et al, 2012_41_, using 0.22NA silica-core multimodal fiber (ThorLabs) epoxied into conical ceramic ferrules of 6.4mm length and 127-131 um bore (Precision Fiber Products, Chula Vista, CA). Fibers were polished using lapping sheets and tested using the LED system and an optical power meter to monitor output (OptoEngine PSU-III OptoEngine LLC, Midvale, UT and ThorLabs PM100D). Fibers with <80% power transmission or poorly defined, non-concentric light output were discarded. Optical fibers were implanted into male C57Bl/6 mice vertically, with the tip of the fiber targeting hippocampal efferent projections upstream of the BNST, with coordinates (mm to Bregma: 0 AP, ±1.2 Lat, 3.8 D/V). Optical fibers were held in position using headcaps constructed of Den-mat dual-cure Geristore two-part dental cement (DenMat Holdings LLC, Ref 4506 and 03452410, Lompoc, CA). The skull was cleaned thoroughly and prepared with Vetbond (3M, Maplewood, Minnesota) to reduce skull moisture and promote skullcap adhesion. Fiber implants left a small but noticeable disruption of brain tissue that was apparent following fixation of brain, and this disruption was used for verification of fiber placement. After the completion of the experiment, animals were euthanized, and the brains were prepared for imaging as described in section 2.3. Following visual analysis, animals without both accurately targeted and high levels of hippocampal viral expression and appropriate fiber placement were excluded from the results (n = 2/9, 1/9 mice respectively). All imaging verification was done blinded to the experimental results.

Five weeks were allowed following surgery for mice to recover and for the virus to fully express and transport ChR to distal axon terminals. Animals were singly housed two days prior to, and for the duration of, the experiment to minimize the stress of changing cage environment during the experiment as a source of variability. Animals were randomly assigned to either receive stimulation or mock stimulation on the first trial and then received the inverse treatment during their second trial, with at least one week separating trials in individual animals. All experiments were completed between 7:00 am and 10:00 am (0-3 Zeitgeber time) in order to reduce variability in circulating corticosterone (CORT) at baseline as well as minimizing baseline CORT levels ^42–45^.

On the day of the experiment, all setup of the optical stimulation and blood collection materials were prepared and made ready for use prior to any animal handling. Immediately prior to the stimulation session, the singly housed animal cages were quickly and carefully moved from their housing location to the experimental room. Animals were rapidly withdrawn from their cage and immediately placed into the restraint device. The optical fibers were coupled to the light source and photostimulation and the timing of the restraint began. 1 ms pulses of light stimulation were given at 20Hz in 2 second intervals alternating between stimulation and no stimulation for 15 minutes. 10mW power at the implanted fiber tip was targeted based on measured power output of the light source on the day of the experiment and the fiber power percent transmission recorded prior to implantation ^46^. The initial tail snip and blood was collected <3 minutes after initial handling of the animal’s cage and used for baseline CORT measurements. Further blood collection was performed at 15, 30, 60, and 120 mins after the initial collection.

### 2.6 HPA axis responsivity

Plasma corticosterone was measured following an acute 15 min restraint. Testing occurred 0-3 h after lights on. Tail blood from adult mice (*n* = 6 per condition) was collected at onset and completion of restraint (0 and 15 min, respectively) and 15, 45, and 75 min after the end of restraint (30, 60, and 90 min, respectively). Tail blood collection requires <30 s to complete. 5ul of blood was pipetted into 10ul of 50 mM EDTA buffer immediately following collection. Tubes were then centrifuged at 5000 rpm and plasma was stored at -80°C until RIA analysis. Corticosterone levels were determined by ImmuChem Double Antibody Corticosterone ^125^I-corticosterone Radioimmunoassay Kit for Rats and Mice according to kit instructions (MP Biomedicals, Santa Ana CA). A standard curve was generated for each run of RIA for 0-1000ng/ml concentration, and ^125^I counts were converted to CORT concentrations based on these curves. Samples were run in triplicate, with the average being used. Samples in which the raw counts from the gamma counter differed by >1000 from replicate samples were discarded, and the average of the two remaining samples was used (7/270). Triplicates were each pulled from a single blood collection sample before RIA processing, so poor tail blood samples could lead to incorrect data. Therefore all cumulative CORT data from RIA was subjected to RObust regression and OUtlier removal (ROUT, Graphpad9) prior to analysis, and 2/60 samples were identified as outliers and excluded ^47^.

### 2.7 Statistics

Mixed-effects ANOVA and paired and unpaired t-tests were used to compare mean values between treatment groups where appropriate. For estimating CORT levels in control and treatment group over time, we performed a 2-way mixed method ANOVA, as the samples were taken as repeated measures but incomplete following outlier removal. Area under the curve calculations for the RIA data were performed using trapezoidal calculation based on the time between each sample and the calculated CORT value. Statistics were computed in Prism 9 (Graphpad, San Diego, CA). All data were analyzed while blinded to experimental conditions.

## 3. Results

### 3.1 Distribution of hippocampal axon terminals

To identify potential brain regions that could function as an inhibitory relay for hippocampal regulation of responses, we first identified brain regions that 1) have high densities of GABAergic cells, 2) received excitatory inputs from the vHip, and 3) have known projections to PVN. Anatomical visualizations of ventral hippocampal projections was performed by expressing ChR and EYFP in the ventral hippocampus in mice expressing the red fluorescent protein tdTomato in GAD-cre expressing interneurons (Figure 1a, parasagittal section). Axonal projections from the ventral hippocampus were observed in the fornix, and dense terminal labeling was seen surrounding the anterior commissure within the bed nucleus of the stria terminalis (BNST) (Figure 1b,c,d parasagittal sections). Similar projections were observed in the neuropil surrounding the PVN in the peri-PVN using the same ChR and EYFP injection scheme in a second line of mice expressing tdTomato in CRF-cre expressing neurons. There was little evidence of projections directly to the PVN itself (Figure 1e, coronal section). Expression was primarily seen in the medial portions of the BNST, with some expression anterior to the anterior commissure, and more pronounced expression inferior and posterior to the anterior commissure. These anatomical findings led us to further interrogate the BNST and peri-PVN as potential relays between hippocampus and PVN.

### 3.2 Physiological responses to activation of hippocampal axonal terminals

Using the same injection scheme of ChR-EYFP into the ventral hippocampus, we measured optically evoked responses in CRF+ neurons in the PVN. Coronal sectioning provided planes containing both EYFP-labeled terminals and tdTomato-expressing cells, indicating the presence of hippocampal terminals adjacent to cell bodies in the PVN in the peri-PVN. Cells were recorded using whole-cell voltage-clamp at holding potentials of 0 mV, the reversal potential for glutamatergic currents, and -60mV, the reversal potential for GABA_A_ receptor mediated currents, to isolate optically evoked inhibitory postsynaptic currents (oIPSCs) and excitatory postsynaptic currents (oEPSCs), respectively. Optical stimulation was targeted at regions of dense EYFP expression near the fornix (Supplemental Figure 1a,b). Stimulation of terminals in both peri-PVN and fornix in coronal sections failed to evoke any synaptic currents in CRF+ cells at either potential (18 cells in slices from 6 animals). The failure to detect synaptic currents may have been because the axons were severed. We therefore repeated the experiments in brain sections prepared in the parasagittal plane, as shown in Boudaba et al 1996 ^17^. In these parasagittal slices, oEPSCs and oIPSCs were recorded in CRF+ neurons in response to photostimuli delivered to EYFP-positive terminals in the region of the fornix (Figure 2a), suggesting that these axonal projection systems lie in a predominately anterior-to-posterior orientation that remains intact during parasagittal sectioning. As inhibition from the peri-PVN should remain intact in coronal sections whereas inhibitory projections from the BNST are more likely to be preserved in parasagittal sections, we suggest that hippocampal regulation of the PVN was more likely to be mediated by interposed neurons in the BNST and it was therefore the focus of our further experiments.

**Figure 2.**
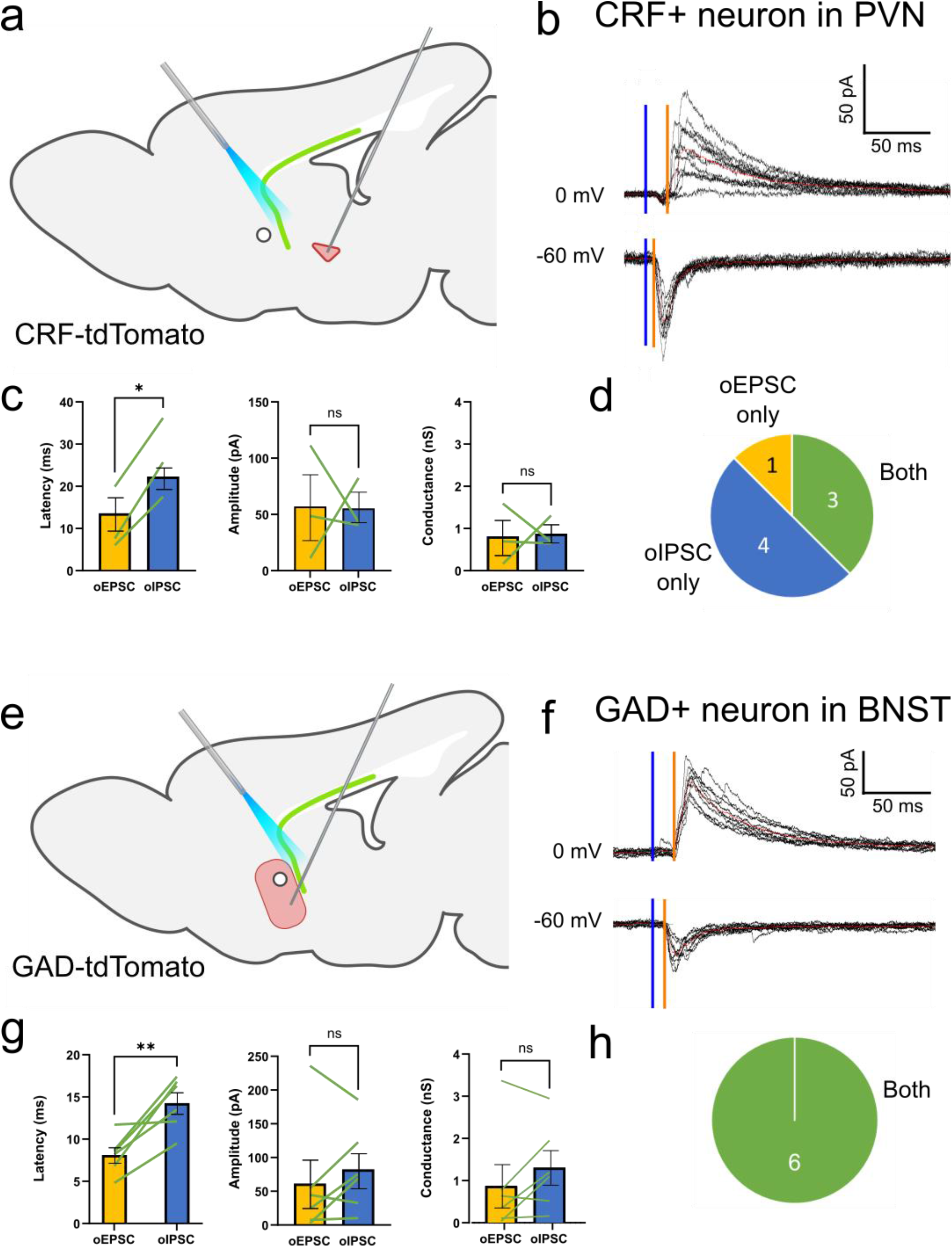
Optically-evoked currents in CRF+ neurons of PVN and GAD+ neurons of the BNST. **a)** ChR and EYFP were expressed in CA1 pyramidal cells of the ventral hippocampus/subiculum in CRF-tdTomato reporter mice (male and female). **b)** Whole-cell voltage-clamp recordings from a CRF+ neurons in the PVN. Photostimulation in the fornix elicited oEPSCs and oIPSCs at holding potentials of -60mV and 0mV, respectively. Sample traces obtained with 10 stimuli delivered 1 minute apart. Optical stimulation delivered at time indicated by blue bar and the latency to oIPSC (0mV) and oEPSC (−60mV) onset shown with orange bar. **c)** Pairwise comparison of oEPSCs and oIPSCs in single cells demonstrate an increased stimulus-response latency for oIPSCs compared to oEPSCs (n=3 pairs, p=.0162), but not significant differences in oEPSC and oIPSC amplitude or conductance. **d)** Number of cells in which oEPSCs, oIPSCs, or both were elicited. **e)** ChR and EYFP were expressed in CA1 pyramidal cells in GAD-tdTomato mice. **f)** Whole-cell voltage-clamp recordings from a GAD+ inhibitory neuron in the BNST. Photostimulation in the fornix elicited oEPSCs and oIPSCs at holding potentials of -60mV and 0mV, respectively. Optical stimulation delivered at time indicated by blue bar and the latency to oIPSC (0mV) and oEPSC (−60mV) onset shown with orange bar. **g)** There was an increased stimulus-response latency for oIPSCs compared to oEPSCs (n=6, p=.0074), but no significant differences in oEPSC and oIPSC amplitude or conductance. **h)** All six BNST cells recorded demonstrated both oEPSCs and oIPSCs. *, p<0.05; **, p<0.01.

Within the PVN, stimulation of hippocampal fibers in the fornix elicited synaptic currents in CRF+ neurons that were dominated by inhibition (Figure 2a-d). The fraction of recordings in which an oIPSC was elicited (7 of 8 cells) was greater than the fraction in which an oEPSC was elicited (4 of 8 cells) (Figure 2d). In experiments in which photostimulation elicited both oEPSCs and oIPSCs (3 of 8 cells), the mean amplitude and conductance of oIPSCs were not significantly different than for oEPSCs (Figure 2c), although statistical comparisons are limited by the low number of cells with both. The IPSCs occurred with a significantly longer latency than EPSCs after photostimulation (Figure 2b,c), and displayed considerable jitter and fractionation, suggesting that the excitation is monosynaptic whereas the inhibition is disynaptic.

We next recorded synaptic responses from GABAergic cells within the BNST. Photostimulation of hippocampal terminals elicited both oEPSCs and oIPSCs in GAD+ cells (Figure 2e-h). In contrast to the PVN, both oIPSCs and oEPSCs were elicited within the same cell in all 6 experiments (Figure 2f,h). Furthermore, the mean amplitude and conductance of evoked EPSCs and IPSCs were comparable in the BNST (Figure 2g). oIPSCs also had a longer latency than oEPSCs in the BNST, suggestive of monosynaptic excitatory stimulation and disynaptic inhibition, presumably by local inhibitory circuits within the BNST ^24^.

### 3.3. High frequency electrical stimulation

Activity in hippocampal inputs to the BNST and PVN *in vivo* is likely to occur at fairly high frequencies, and excitatory and inhibitory synaptic transmission display well-known activity-dependent dynamics ^48,49^. We therefore asked whether the net balance between excitation and inhibition was changed in a frequency-dependent manner in the PVN. Unlike optical stimulation, electrical stimulation of hippocampal afferents in the fornix produced both EPSCs and IPSCs in the majority of cells recorded from (11 of 12) and was thus better suited to compare activity-dependent dynamics. We placed an electrode in the fornix of parasagittal brain slices and recorded electrically evoked responses in CRF+ cells in the PVN using 20 Hz stimulation for 1 second bouts, repeated 10 times. eEPSCs and eIPSCs persisted with 20Hz stimulation, but both gradually became decreased in amplitude and conductance by about 50% over the course of the stimulus train in a roughly parallel fashion (Figure 3a,b). In addition, the percentage of stimuli that failed to elicit responses increased during the train (Figure 3c). There was no significant change in the balance of excitation and inhibition, measured as either conductance or failure rate, during the course of the 20Hz frequency trains, despite the decrease in amplitude.

**Figure 3.**
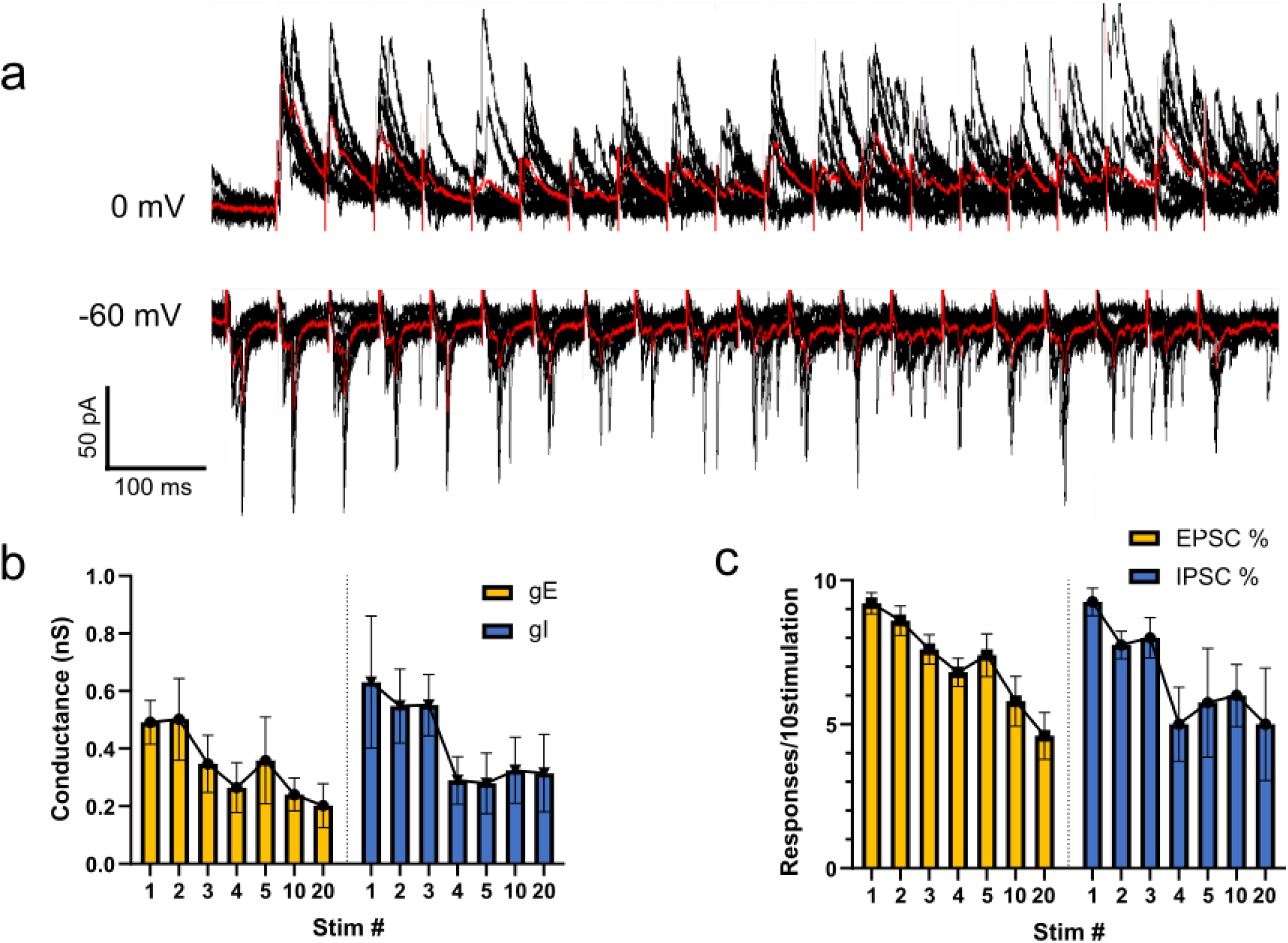
High frequency electrical stimulation of fornix elicits currents in CRF+ neurons in the PVN. **a)** Electrically evoked EPSCs (−60mV) and IPSCs (0mV) recorded in CRF+ neuron in the PVN in response to stimulation of the fornix at 20Hz for 1 sec, repeated 10 times. **b)** There was a comparable ∼50% decrease in both EPSC and IPSC conductance over the duration of the stimulus train. **c)** Similarly, there was a comparable ∼50% decrease in the probability that a given stimulus elicited an EPSC or IPSC over the duration of the 20Hz stimulus trains.

### 3.4. Effects of hippocampal output on HPA axis function in vivo

Previous studies have shown that lesions of GABAergic cells in the BNST lead to an enhancement of HPA axis responses to acute restraint stress^23^, but it has never been shown that hippocampal activation acutely regulates the stress-response. Because our anatomical and electrophysiological data suggested that the hippocampus inhibited CRF+ PVN neurons via activation of the BNST, we predicted that stimulating hippocampal afferents to the BNST and PVN would attenuate the plasma glucocorticoid response to an acute stressor. We expressed ChR-EYFP bilaterally the ventral hippocampus of wildtype C57/Bl6 mice, as above, and implanted two optical fibers with the fiber tips targeting the fornix region upstream of the BNST and PVN. Mice were placed in a restraint tube and connected to dual optical fibers. After 15 minutes, they were returned to their home cages and the fiber was detached. Blood was collected via tail snip at 0, 15, 30, 60, and 120 minutes after initial placement in the restraint tube. As a control for variations in ChR expression and fiber placement, a cross-over experimental design was used, in which mice received optical stimulation at 20Hz frequency 10 mW intensity in an alternating 2s on/off stimulation protocol for the duration of the 15-minute acute restraint stress, or a mock-stimulation trial (Figure 4a). The mock stimulation included all the handling and light associated with the real stimulation, but the optical pathway at the ferrule sleeve was blocked with lens tissue, reducing the measured output intensity by more than 100-fold to <0.1 mW. The restraint and stimulation procedure were performed in two separate trials one week apart for each animal to generate within-animal paired datasets (Figure 4a). Acute restraint stress produced a transient elevation of plasma CORT level that peaked 30-60 min after the onset of stress and declined over the following two hours (Figure 4b-d). Mice receiving photostimulation at 10 mW had a significant decrease in the peak stress-induced CORT levels, compared to the same animals receiving mock stimulation (Figure 4b-d, Suppl. Figure 2a,b). The effects of stimulation at 10mW persisted beyond the period of stimulation, as apparent in different total areas under the curve (Figure 4c).

**Figure 4.**
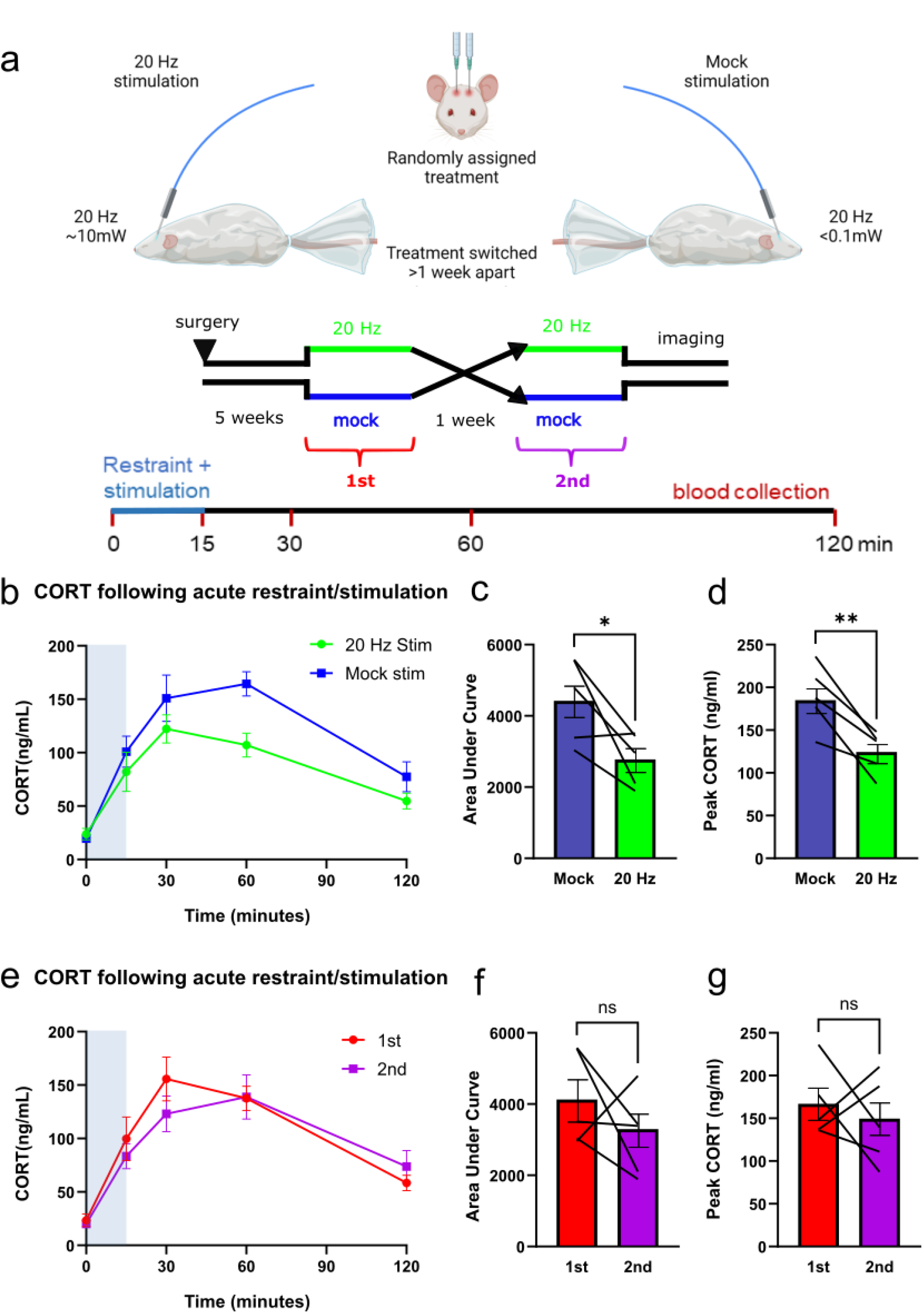
Optical stimulation of hippocampal efferents suppresses the stress-induced CORT response. **a)** ChR and EYFP were expressed in the CA1 region of male C57/Bl6 mice bilaterally and optical fibers targeted hippocampal terminals in the fornix anterior to the BNST. Each animal was randomly assigned to 20Hz stimulation at 10mW or mock stimulation at 0.1 mW and the opposite treatment 1 week later. Animals were restrained for 15 minutes and received 20Hz optical stimulation 2 seconds on, 2 seconds off for the duration of the restraint. Blood was sampled via tail snip at 0, 15, 30, 60, and 120 minute timepoints for CORT assay. **b)** Comparison of the timecourse of CORT responses in the full power and mock 20Hz stimulation trials. Mixed-effects ANOVA was performed on the resulting curve, with time effect (n=6, p<0.0001) and treatment effects (p=0.0460). **c**,**d)** Comparison of the area under the curve and peak CORT responses in the paired trials from b showing significant effects of hippocampal stimulation. Paired t-tests for mock vs stim AUC (p=0.0184) and peak value (p=0.0105) were significant. *, p<0.05; **, p<0.01. **e**,**f**,**g)** The same data after resorting by recording order demonstrating no significant effects. When segregated by order of recording, mixed effects ANOVA maintains time effect (p<.0001) but loses treatment effect (p=0.5333). Similarly, recording order AUC (p=0.2936) and peak CORT (p=0.5404) were not significantly different.

To control for stimulation order, the CORT values for these mice were regrouped into whether the samples taken during the first or second trial of each mouse (Fig 4a). No significant effect of stimulation order were observed when the data was reanalyzed in this manner (Figure 4e-g, Suppl. Figure 2c,d), suggesting that the strong photostimulation itself was cause for the decrease in CORT. We conclude that hippocampal output is sufficient to partially suppress acute HPA axis responses to stress.

## 4. Discussion

The synaptic mechanisms underlying hippocampal regulation of the HPA axis were investigated in mice. Because the hippocampus is known to be both stress-sensitive and associated with psychiatric illness, understanding its contributions to neuroendocrine regulation is important for many disease processes. The canonical hypothesis is that the hippocampus decreases stress-induced glucocorticoid production via inhibition of the CRF-releasing cells in the PVN. The major projection neurons of the hippocampus are excitatory, therefore, for this hypothesis to be true there must be one or more inhibitory brain regions interposed between the hippocampus and the PVN. The ventral hippocampus is considered to be the region of the hippocampus most responsible for processing affective information, particularly in regards to stress and anxiety and therefore was the focus of this study ^12,13,50,51^.

Prior studies have suggested that the hippocampus inhibits production of glucocorticoids, and the high expression of glucocorticoid receptors in the hippocampus make it likely to be involved in negative feedback of the HPA axis. Although this idea is largely accepted, the mechanisms and neural circuits underlying this response are unclear and to our knowledge there is no direct evidence of the effects of hippocampal projections on CRF-releasing cells in the PVN. It was also uncertain if hippocampal activity is even capable of inhibiting the HPA axis during acute stressors, or if it takes more a sustained and chronic perturbation in hippocampal activity to meaningfully alter the HPA axis.

Consistent with prior anatomical studies, observation of hippocampal nerve terminals labeled with virally expressed EYFP demonstrated abundant innervation of the BNST and peri-PVN region, which were previously identified as likely targets for this regulation ^23,52^. There were few hippocampal projections directly innervating the regions of the PVN where CRF+ cell somata are located, also consistent with previous work ^23^. It is noteworthy that a similar paucity of direct excitatory projections to the PVN, but strong projections to the BNST, has also been reported for efferents from the infralimbic (IL) cortex ^53^, suggesting that they produce similar effects on HPA axis function.

We used optogenetic stimulation of hippocampal efferents in transgenic reporter mice to test for synaptic responses in CRF+ neurons in the PVN. Although we could clearly see both terminals and CRF+ cells in the PVN within a single slice cut in the coronal plane, we were unable to optically evoke responses. In parasagittal sections, in contrast, synaptic responses were consistently evoked with the identical stimulation and recording configuration. We suggest that this may explain why these responses have not been previously described in the literature. Boudaba et al 1996 ^17^ did evoke inhibitory responses in PVN cells in response to focal application of glutamate in the peri-PVN in coronal slices, suggesting that the GABAergic cells and their projections to the PVN remain intact in such slices. Optical stimulation of hippocampal terminals in the peri-PVN failed to produce an equivalent response, however, suggesting that hippocampus does not inhibit the PVN via activation of these peri-PVN inhibitory neurons.

With optical stimulation in parasagittal sections, we were twice as likely to elicit inhibitory responses in CRF+ cells as excitatory responses. Although there is little discussion of direct excitatory input from hippocampus to the PVN in prior literature, we did record optically evoked EPSCs in 4 of 8 recordings. In 3 of 8 cells, both excitatory and inhibitory responses were optically evoked. The latency for IPSCs was twice as long as for EPSCs, indicative of a direct excitatory input and indirect disynaptic or polysynaptic circuit for the inhibitory inputs.

In contrast to the responses elicited with selective optical stimulation of hippocampal afferents, electrical stimulation within the fornix elicited both EPSCs and IPSCs in 11 of 12 cells. We suggest that this difference results because electrical stimulation recruits some other, stronger direct excitatory input to the CRF+ cells in the PVN, such as from the amygdala, rather than just ventral hippocampal inputs.^54^ There was no evidence of a frequency-dependent shift in the balance between excitation and inhibition with electrical stimulation, suggesting that this does not explain the net inhibitory effect of hippocampal output. Instead, the larger proportion of CRF+ cells displaying inhibitory responses to stimulation of hippocampal efferents is consistent with a potent, divergent inhibitory input from the BNST and perhaps other nearby regions. This strong disynaptic input may be sufficient to explain a net inhibitory effect of hippocampal inputs to the PVN. Another possibility is that the hippocampus has multiple output streams that can alternatively activate an excitatory or inhibitory pathway to the PVN depending on the context of the stressor. There is some evidence of differential effects of subicular lesions depending on the stressor type, suggesting it may differentially play an excitatory or inhibitory role ^55^.

The BNST has long been implicated as a key sign-reversing node in which excitatory output from the hippocampus and IL cortex is converted to inhibition of the PVN ^15,23,56–58^. As predicted, we observed strong excitatory and feedback inhibitory synaptic responses in GAD+ inhibitory neurons within the BNST in response to optical stimulation of hippocampal efferents. Unlike responses in the PVN, optically evoked EPSCs and IPSCs were reliably observed in BNST neurons (7 of 7 cells) and had relatively equivalent conductances. Latency from the optical stimulus was longer for IPSCs than EPSCs, consistent with disynaptic inhibition, either feedforward or feedback. The posterior BNST is reported to inhibit PVN activity ^28^ and our data demonstrates that these cells, as well as some medial regions of the BNST, receive potent excitation from the ventral hippocampus. Description of BNST subregions in the sagittal plane is challenging because most descriptions are in the coronal plane. However, areas immediately posterior or inferior to the anterior commissure, in very medial parasagittal to midline sections, were the most reliable areas for recording evoked responses.

The disynaptic, sign-reversing circuit characterized above provides the potential means for the hippocampus to inhibit the HPA axis, as long postulated, although this has not been previously demonstrated. We observed that both EPSC and IPSC amplitude declined during maintained 20 Hz stimulation but were not eliminated. Furthermore, the balance of excitatory-to-inhibitory conductance remained relatively unaffected by 20 Hz stimulation. These data suggest that sustained hippocampal output would remain largely inhibitory in PVN CRF+ cells.

Using 15 min of restraint as an acute stressor, we observed that simultaneous delivery of 20 Hz optical stimulation of hippocampal afferents to the BNST region produced a significant reduction in the peak and overall elevation of circulating CORT in male mice. This effect was apparent at the cessation of the stimulation and persisted during the full 2 hours of recovery. The consistent low CORT levels at timepoint 0 (<50ng/ml) in both treatment conditions indicates that the mice remained unstressed prior to being placed in the restraint tubes. To our knowledge, this is the first demonstration that stimulation of hippocampal inputs to PVN leads to inhibition of stress-induced HPA axis activation.

Taken together with the complimentary findings that lesions of GABAergic cells within the BNST lead to increased stress-induced HPA activation ^23^, we suggest that the inhibitory projection from the BNST to the PVN accounts for this inhibition. Because the BNST is considered an integrator of input from many brain regions, it was not clear if hippocampus alone would be able to alter systemic levels of CORT. The effect of 20 Hz stimulation was not large, representing a <ca. 25% reduction in the full area under the curve. This suggests that there are other powerful excitatory influences on the CRF cells that the hippocampal-BNST mediated inhibition cannot fully counteract. The effect of optogenetic stimulation of IL cortical efferents produced a comparably sized effect on stress-induced CORT levels ^59^.

These findings suggest that the role of hippocampal projections to the PVN is perhaps not to produce rapid corticosterone-dependent feedback inhibition in response to elevations in CORT levels, as has long been assumed. Indeed, GRs and MRs in the hippocampus are poorly suited to mediate rapid responses and changes in hippocampal output, given the slow nature of the expected responses they produce. Instead, ongoing activity in hippocampal and IL cortical inputs may set an allostatic steady-state tone in the average level of excitability of PVN CRF-releasing cells. These inputs could also trigger tonic changes in PVN tone in response to environmental conditions and features perceived through cortical processing, such as those that might demand a heightened or dampened HPA axis response. More information about the dynamics of hippocampal output to the hypothalamus in response to stress would be valuable in testing these hypotheses.

Optogenetic stimulation of IL efferents to the hypothalamus is inhibitory only in male rats, and not in females ^60^. We note that our experiments were only performed in male mice because we were unable to find a description of the anatomy of the BNST in female mice to guide placement of the optical fiber. Literature suggests that the female BNST is several fold smaller in volume than the male BNST ^20,61^, but we found no reliable source demonstrating where these volumetric changes are concentrated. There is also a potential sex-difference in the hippocampal input to PVN as well as the multitude of other brain regions that signal to the BNST ^62^.

The hippocampus has numerous links to both major depression and neuroendocrine regulation in clinical and preclinical literature ^63–65^. Using rodent models, as we and others have shown there are lasting anatomical and physiological changes in hippocampal synapses following chronic stress that correlate with depression-relevant behaviors ^66,67^. These synaptic changes are reversible following administration of antidepressant compounds and resolve with the behavioral measures ^67,68^. Furthermore, treatment with the glucocorticoid corticosterone (CORT) was shown to mimic the behavioral and synaptic changes seen following chronic stress ^9^. Blocking endogenous production of CORT during chronic stress was able to block the stress-induced synaptic and behavioral changes. We and others have also previously shown functional significance of hippocampal projections in mediating reward-related behavior ^69–71^. Our present findings suggest that dysregulation of the HPA axis, a symptom of depression, may also involve similar hippocampal mechanisms. Volumetric changes in the hippocampus are seen in patients with major depression, and lesion studies demonstrate that loss of hippocampus or the major output of hippocampus can lead to dysregulation of the neuroendocrine system ^14,72,73^. The ventral hippocampus is of particular interest, as studies show increased connectivity to affective brain regions (for review, ^51^). It will be of interest to investigate the possible role of changes in hippocampal inputs to the PVN and BNST under conditions of stress-response dysregulation such, as is seen following chronic stress.

## 5. Conclusion

We have shown that hippocampal afferents elicit monosynaptic excitation and disynaptic inhibition of CRF-secreting cells in the PVN and that activation of this projection can significantly alter the acute stress response. Further understanding the central regulation of PVN in both normal and disease states is essential and could lead to important breakthroughs in our understanding of disease and improving treatment options.

## Acknowledgements

This work was supported by NIH grants R01 MH086828 (SMT); MH108286 and ES028202 (TLB); T32 GM092237 to the Medical Scientist Training Program; T32 GM008181-30 Training Program in Integrative Membrane Biology (ABC); and S10 OD026698 to the University of Maryland Microscopy Core Facility.

We thank the following for assistance in this manuscript:

Andreas Wulff for insightful comments during the writing and editing process, as well as assistance in creation of figures.

Dr. Joseph Mauban and the University of Maryland School of Medicine Center for Innovative Biomedical Resources Microscopy Core Facility for training and use of the W1 spinning disc confocal microscope.

Andrea Romanowski and Dr. Alexandros Poulopoulos at the University of Maryland Department of Pharmacology for assistance and use of equipment in imaging using the Ti2 epifluorescence microscope.

Dr. Tara Legates for training and encouragement in patch-clamp electrophysiology and optogenetics.

Dr. Kathleen Morrison for assistance in radioimmunoassay training and CORT response expertise.

Figure 2 and Figure 4 were created in part using BioRender.com.

## Figure legends

**Supplemental Figure 1.**
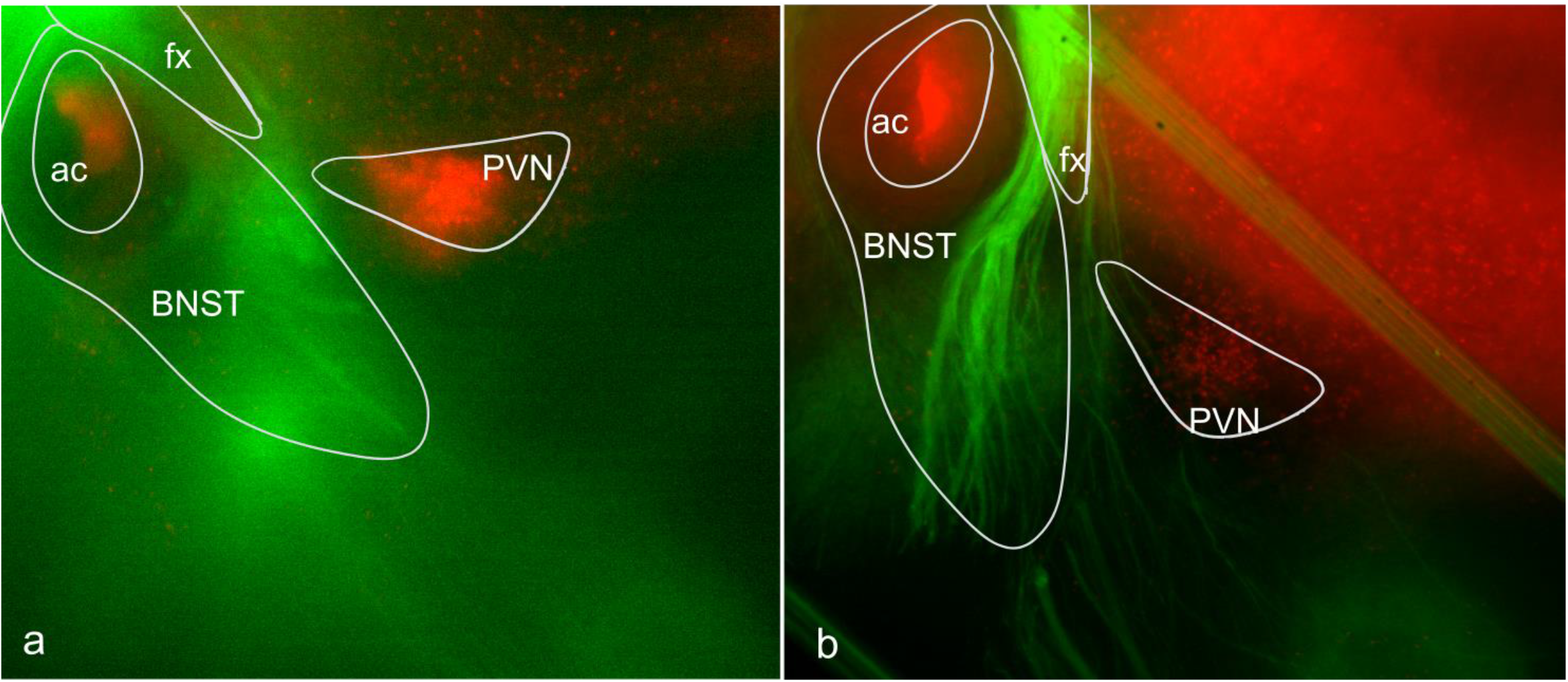
Recording configuration in CRF-tdTomato mouse brain slices. Two widefield images taken on the recording setup with CRF+ neurons (red) in CRF-tdTomato reporter mice and hippocampal fibers (green) entering the BNST and PVN in parasagittal sections. Hippocampal fibers are dense within the BNST and sparse in the PVN. fx – fornix, ac – anterior commissure, BNST – bed nucleus of the stria terminalis, PVN – paraventricular nucleus of the hypothalamus. Note that the harp used to hold the slice down in the recording chamber causes non-specific fluorescence in the top right portion of b.

